# Identification of phenotypic and transcriptomic signatures underpinning maize crown root systems

**DOI:** 10.1101/2024.08.20.608626

**Authors:** Jodi B. Callwood, Ella G. Townsend, Shikha Malik, Melissa A. Draves, Jasper Khor, Jackson P. Marshall, Heather Sweers, Craig L. Cowling, Justin W. Walley, Dior R. Kelley

## Abstract

Maize is pivotal in supporting global agriculture and addressing food security challenges. Crop root systems are critical for water uptake and nutrient acquisition, which impacts yield. Quantitative trait phenotyping is essential to understand better the genetic factors underpinning maize root growth and development. Root systems are challenging to phenotype given their below-ground, soil-bound nature. In addition, manual trait annotations of root images are tedious and can lead to inaccuracies and inconsistencies between individuals, resulting in data discrepancies. To address these issues, we have developed an automated phenotyping pipeline for field-grown maize crown roots by leveraging open-source software. Phenotypic variation of 20 maize genotypes from the Wisconsin Diversity panel was significant for numerous root traits, suggesting a genetic basis for the observed developmental deviations. In addition, juvenile root traits from controlled environment conditions exhibited inconsistent correlation with field-grown adult root traits, underscoring the developmental plasticity prevalent during maize root morphogenesis. Transcripts involved in hormone signaling and stress responses were among differentially expressed genes in roots from 20 maize genotypes, suggesting many molecular processes may underlie the observed phenotypic variance. This study furthers our understanding of genotype-phenotype relationships, which is relevant for informing agricultural strategies to improve maize root physiology.

## Introduction

Roots are an essential plant organ for water and nutrient uptake (Steudle 2000). Because roots are typically in soil and hidden from the eye’s view, it is challenging to study roots *in situ.* The development of transparent rhizoboxes and rhizotrons has enabled imaging of root systems within soil matrices (Schmidt, Lowry, and Gaudin 2018; Mohamed et al. 2017) and rolled towel assays allow observation of seedling root systems (Draves et al. 2022; He et al. 2023) . However, these approaches can be cumbersome and expensive (Gärtner et al. 2009; Chen, Djalovic, and Rengel 2015; Guo et al. 2013). To better understand crop root growth and development, alternative strategies for phenotyping root systems are required.

*Zea mays* (maize) root systems provide anchorage and exhibit unique phenotypes that aid in plant physiology (Vogt et al. 1995). Genetic and phenotypic differences in maize root architecture are well documented, and ideal root phenotypes for different environmental challenges have been proposed (P. Li et al. 2024). Identifiable root phenotypes include root number, length, surface area, diameter, root angle, average diameter, network area, and angle frequency (Seethepalli et al. 2021a). These root phenotypes were detected through RhizoVision Explorer (Seethepalli et al. 2021a) +, and are valuable in exploring the relationships between root traits and environmental factors (Paez-Garcia et al. 2015). For example, root length correlates to soil physical attributes and lateral root number corresponds to soil nutrients (Amtmann, Bennett, and Henry 2022). Root architecture corresponds to water and nutrient availability within the soil; much can be gleaned about the concentration of nutrients and water resources accessible to a plant through its root architecture. Root branching correlates to nitrogen availability within the soil, roots’ connection to water, and nutrient acquisition leads to unique architecture to survive in the available concentrations of nutrients available. The quantification of traits such as root angle, diameter, quantity, and length can provide insight into the impact of available nutrients on crop development.

Root phenotyping continues to be a significant bottleneck in data analysis. Several root phenotyping pipelines have been developed for these purposes to ease this burden, with various advantages and disadvantages depending on the complexity of the root structure and source images. Root phenotype extraction through 2D images of crown root systems has been accomplished using several different software packages, including EZ-Rhizo (Armengaud et al. 2009), RootNav (Pound et al. 2013), ARIA (Pace et al. 2014), DIRT (S. Liu et al. 2021; Das et al. 2015), and RhizoVision Explorer (Seethepalli et al. 2021a). ImageJ/Fiji allows manual measurement of images, scale adjustments, and conversion from pixels to centimeters. While there is a low threshold to user adaptation of ImageJ/Fiji this software is tedious and low throughput (Schneider, Rasband, and Eliceiri 2012). ARIA 2D allows image analysis of root phenotypes, such as total root length, primary root length, secondary root length, perimeter, center of mass, total number of roots, convex area, network area, bushiness, SLR, center of point, and many others, but it does not account for hole correction for highly overlapping root architecture and relies on Mathematica (Pace et al. 2014). The RootNav 2.0 segmentation platform has been used for semi-automatic root phenotyping that can be adapted to new datasets however the interface does not allow for hyper tuning and the interface is cumbersome also datatype RSML is not universal usage (Yasrab et al. 2019). DIRT an online platform for high-throughput phenotyping of 2D images with many supporting extensions for various root imaging tasks and analysis methods (S. Liu et al. 2021). PlantCV is a comprehensive and customizable image analysis platform for many plant phenotypes, including roots, but it has limited thresholding capabilities and does not support high noise in images (Gehan et al. 2017). RhizoVision Explorer provides a high-throughput 2D image phenotype extraction platform for roots, but the segmentation is not optimized for files with a variation of image topography (Seethepalli et al. 2021a).

Convolutional neural networks (CNNs) are computer vision model architectures that have shaped plant image analysis (French et al. 2009; Cai et al. 2015; Gage et al. 2017). RootPainter is a CNN trainable through a user-friendly graphical user interface (GUI) (Smith et al., 2022). RootPainter performs automated image segmentation and can differentiate the background from objects of interest (Smith et al. 2022; Bauer et al. 2022). This supervised learning method allows RootPainter to be trained on various datasets and thus offers a customizable opportunity for high-throughput phenotyping.

In this study, we sought to leverage existing open-source tools to enable root phenomics. For high-throughput image processing of soil-grown maize crown roots, we developed a phenotypic extraction pipeline using a combination of ImageJ/Fiji, RootPainter, and RhizoVision. This workflow efficiently rapidly phenotyped >50 key root traits in 20 maize genotypes, demonstrating the wide phenotypic range of field-grown corn roots. In addition, we compared these data to manually phenotyped juvenile root traits to determine how well early seedling root traits correlate with adult traits. Overall, we observed inconsistent correlation between developmental stages, highlighting the phenotypic plasticity prevalent in most maize genotypes during root morphogenesis. A transcriptomic analysis of seedling root systems indicates a wide range in gene expression patterns related to biological processes including hormone pathways, stress responses, and cell walls. Thus, the wide variance observed in maize root development is likely due to the interplay between many complex pathways. Overall, this approach enables rapid and facile root trait quantification, thus enabling integration of transcriptomic and phenomic datasets to generate new hypotheses about genotype-phenotype relationships across root developmental stages.

## Materials and Methods

### Experimental and Technical Design

An overview of the experimental and technical design performed in this study is grouped by developmental age and type of analyses performed (**Figure 1**). Briefly, mature crown root phenotyping of 22 maize genotypes was carried out using images captured from a DSLR camera that were cropped and trained in RootPainter (Smith et al. 2022). After training, all images were analyzed in RhizoVision. In parallel, juvenile root phenotyping was performed with images of 10-day-old maize seedlings from 11 genotypes grown in rolled towel assays captured with smartphones. Seedling root traits were subsequently hand-annotated and measured using ImageJ/Fiji. Following image acquisition, root tissues were harvested from 5-10 seedlings per genotype and analyzed for transcript abundance using 3’ end RNA sequencing. Differential gene expression analysis and Gene Ontology (GO) enrichment analyses were performed on the transcriptomic data. Finally, a correlation analysis between the adult crown root traits quantified via RhizoVision and manually curated juvenile seedling root traits was carried out using Spearman correlation.

**Figure 1.**
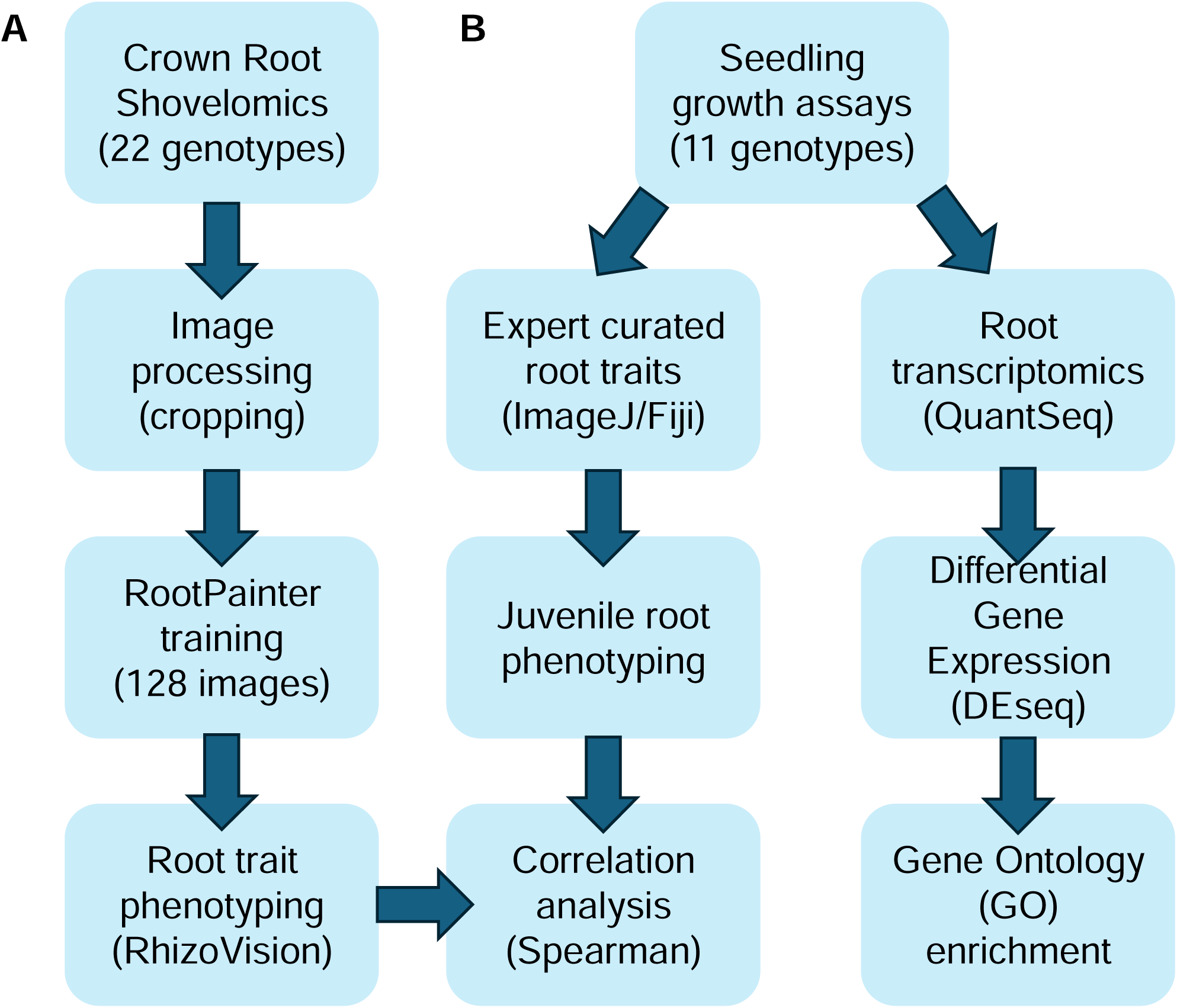
Experimental workflow. (A) Shovelomics was performed on 22 maize genotypes from the Wisconsin Diversity panel to image crown roots at stages V6-V10. Prior to training in RootPainter, images were cropped to remove the background. RootPainter was trained on 128 crown root images to correctly annotate root over the background. Root traits were quantified using RhizoVision. (B) Rolled towel growth assays were performed on 11 genotypes from the Wisconsin Diversity panel to quantify juvenile root traits in 7-day-old seedlings. Primary root length was manually annotated using ImageJ. After imaging seedlings, root tissue was harvested for RNA extraction and transcriptomics via QuantSeq. Differentially expressed genes (DEGs) in the examined genotypes compared to the reference genome, B73, were identified using DEseq. Gene Ontology (GO) enrichment analysis was performed on the DEGs. Correlation between juvenile and adult root traits was assessed using the Spearman coefficient.

### Plant materials

For crown root phenotyping from field-grown maize inbreds, 22 genotypes were selected from the Wisconsin Diversity panel as proof-of-concept (Supplemental Table S1). All genotypes were planted in a random block design at the Iowa State University Agricultural Engineering/Agronomy farm, Boone, IA, in the summer of 2022. Each genotype was grown in twin rows, with ten plants per row, in two separate locations in the same nursery to minimize plot effects. At stage V6-V10, three plants per genotype and row were excavated from the field to generate 12 biological replicates per genotype for image analysis. The range in growth stages was due to the weather and differences in germination rate among the genotypes examined. Harvested plants were taken from the middle of each row to avoid end-of-row effects. Individual plants were tagged with slip-n-lock vinyl labels for identification. All stem material above the third node was removed with Corona Forged steel 8” shears and discarded. Crown root systems were soaked in room temperature water for 10-30 minutes prior to washing with a commercial sink sprayer. Root systems were air dried for six days prior to imaging. Images were acquired with a Canon E0S 80D DSLR camera with an 18-55 lens using a ring light and black vinyl photography background.

For the transcriptomic analysis, ten-day-old maize seedlings were grown using the rolled towel method as previously described (Draves et al. 2022). Root tissue from 6-10 seedlings was harvested and pooled together for each biological replicate, flash-frozen in liquid nitrogen, and stored at -80°C.

### Image analysis

A random subset of 255 images from the crown root image dataset (Kelley et al. 2024) was selected for analysis and training of RootPainter (Smith et al. 2022). When possible, these images were cropped to reduce background noise, including brace roots and stem tissue. This image subset was randomly separated into training (50%), testing (30%), and confirmation (20%). A training set of 128 images was used to train RootPainter to segment root systems from the background. To remove residual noise in the image background or from stem and/or brace root tissues, mean crown root diameters were manually calculated from 271 images using ImageJ (Schneider, Rasband, and Eliceiri 2012). The resulting segmented images were analyzed in RhizoVision (Seethepalli et al. 2021b). Diameter thresholding at eight ranges of 0-200mm, 200-400mm, 400-600mm, 600-800mm, 800-1000mm, 1000-2000mm, 2000-3000mm, and over 3000mm were set in RhizoVision. Segments with thresholds over 3000mm were identified as stem tissue and discarded from the subsequent analysis. The pipeline took approximately seven hours to process all the 235 images. Each trait was normalized within genotype, and mean values across genotypes were compared to B73, the reference genotype (Supplemental Table S1). Statistical significance was assessed using one-way ANOVA in GraphPad Prism.

### RNA extraction, library preparation and next-gen sequencing

Root tissue from 10-day-old seedlings grown via rolled towels ground to a homogenous fine powder using a mortar and pestle. For each inbred, a pooled sample of 5-10 roots per genotype was harvested into a singular biological replicate. For B73, two biological replicates were collected. Total RNA was extracted from ground root samples using TRIzol™ Reagent followed by purification using Zymo RNA MiniPrep kit. RNA quality and concentration was determined by Qubit RNA BR assay. For each genotype, 500 ng of total RNA was used to make QuantSeq libraries using Lexogen QuantSeq 3’ mNRA Library prep kit FWD for Illumina (UDI 12nt). Library construction was measured with the Qubit 1X dsDNA high-sensitivity assay. The library integrity was checked with the AATI fragment analyzer. An average of 310 bp library size was observed. QuantSeq libraries were sequenced on a NovaSeq 6000 with a read length of 100 nt at the ISU DNA facility.

### Transcriptomic analyses

Reads obtained after sequencing were quality-checked using FastQC. Adaptors were trimmed using Cutadapt (cutadapt/2.5-py3-j2ra6iu). Reads were aligned to ZmB73V4 using STAR (STAR_index_2.7.5a_ZmB73V4) and B73_RefGen_v4_genomic.gtf annotation file. Indexing was performed using samtools/1.9-k6deoga and counts were obtained using Featurecounts. In total 31,019 transcripts were detected (Table 1). Differentially expressed genes (DEG) were identified using DESeq2 (Love et al. 2015) by comparing each genotype to B73 with an FDR (adjusted *P* value) of ≤ 0.05 (Table 1; Supplemental Table S2).

**Table 1.** Summary of differentially expressed genes (DEG) in root tissue from 10 maize genotypes compared to B73. Total DEG at false discovery rate (FDR) £0.05. The number of DEG in each genotype after applying a log2 fold change (log2FC) cutoff of ± 1 for up and down-regulated transcripts is shown.

| Genotypes | Total Transcripts DE | log2FC ± 2 | | |
| --- | --- | --- | --- | --- |
|  |  | DEG | Up | Down |
| A239 | 3568 | 2234 | 719 | 1515 |
| MS225 | 3817 | 2286 | 830 | 1456 |
| N201 | 140 | 79 | 25 | 54 |
| ND247 | 3638 | 2206 | 792 | 1414 |
| OS602 | 3352 | 2056 | 774 | 1282 |
| W153R | 4477 | 2436 | 930 | 1506 |
| W604S | 3479 | 2140 | 783 | 1357 |
| W819G | 1831 | 1060 | 327 | 733 |
| W9 | 4061 | 2391 | 979 | 1412 |
| WR3 | 4168 | 2517 | 943 | 1574 |

Differential gene expression was determined by identification of transcripts exhibiting significant alterations in a genotype relative to B73. A log2 fold-change (FC) was used to define increased and decreased transcripts compared to B73. Visualization of the top 100 differentially expressed genes (DEG) was performed by hierarchical clustering implemented via the heatmap package (Kolde, 2019).

A <u>d</u>ensity-<u>b</u>ased <u>s</u>patial <u>c</u>lustering of <u>a</u>pplications with <u>n</u>oise (DBSCAN) algorithm (Mahesh Kumar and Rama Mohan Reddy 2016) cluster plot was generated for each genotype analysis. Approximately two to three clusters of DEGs were detected per genotype from DBSCAN (Supplemental Table S3).

DEG were subjected to Gene Ontology (GO) analysis of biological process (BP) terms (Ge, Jung, and Yao 2020). Terms exhibiting FDR of ≤ 0.05 based on a Fisher’s statistical test and Yekutieli adjustment method were deemed statistically significant (Supplemental Table S4). The background for comparison was all transcripts identified in this experiment.

### Juvenile and Adult Root Trait Correlation Analysis

Spearman correlation analyses were performed to ascertain whether seedling root traits correlate with adult crown root traits (Supplemental Table S5). Normalization through resampling and analysis were done in R using dplyr (Wickham et al. 2023), tidyverse (Wickham et al. 2019), openxlsx (Schauberger and Walker 2022), and base statistics.

## Results

### Pipeline for phenotyping maize crown roots

Maize crown root systems can exhibit a wide range of phenotypic plasticity (Sha et al. 2023; York and Lynch 2015; Hochholdinger, Yu, and Marcon 2018). To enable high throughput analysis of field-grown maize crown root images, 22 inbreds were imaged at V6-V10 with replication. RootPainter was selected for training because it is freely available and can integrate with RhizoVision (Smith et al., 2022; Bauer et al., 2022). After training RootPainter with 128 crown root images, a total of 271 crown root images were processed through the pipeline (**Fig. 1A**). Each image took approximately 1 minute to process using RootPainter. ImageJ was used to clean up any residual background as needed, taking around 30 seconds per image. In total, image preparation for RhizoVision took approximately 1.5 minutes per image. RhizoVision took around 20 minutes to process all images for 56 traits. Processing 271 images took approximately 7 hours (**Fig. 2**).

**Figure 2.**
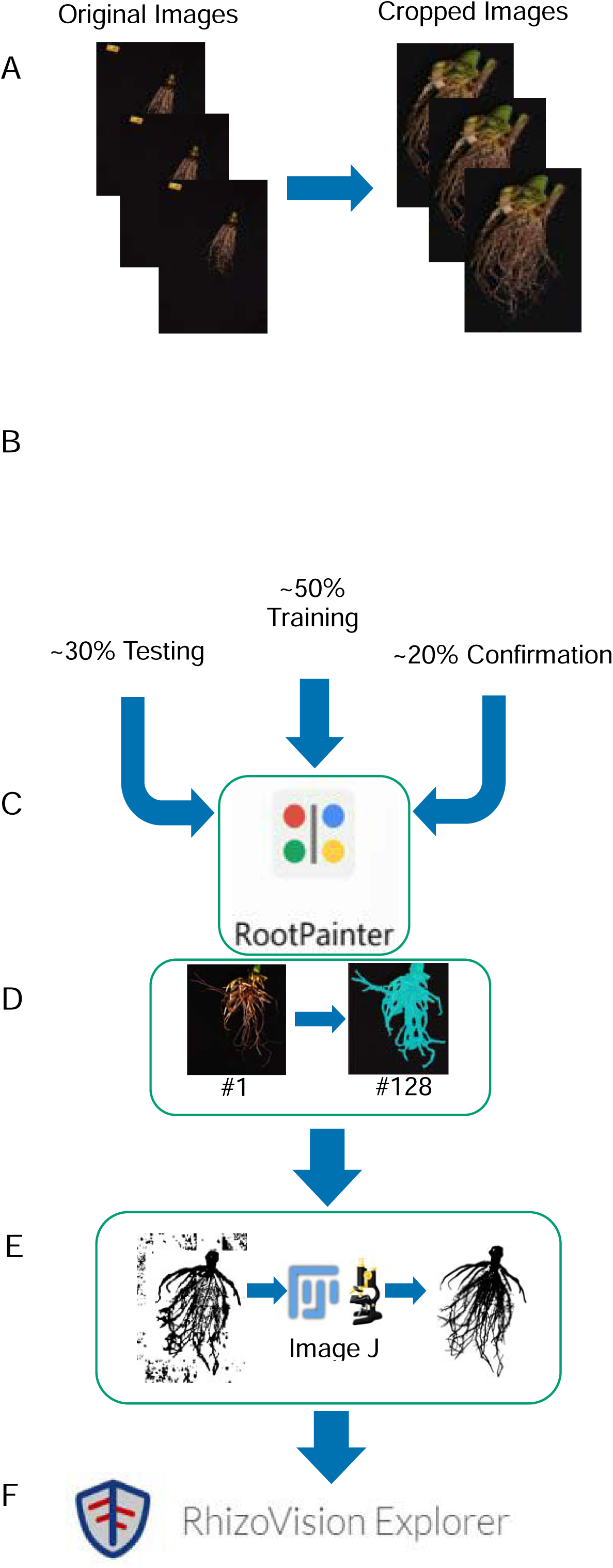
Pipeline of crown root image analysis for trait quantification. (**A**) A subset of mature crown root images was initially cropped to remove the background. (**B**) Cropped image files were randomly split into three groups: training (∼ 50%), testing (∼30%), and confirmation (∼20%). (**C**) The training images were fed into RootPainter, a U-net application, for training. (**D**) Exposing RootPainter to ∼128 images to differentiate between two parameters (root and not-root). These images were used to train RootPainter for segmentation. (**E**) Output images often required editing in ImageJ/Fiji to remove the background. (**F**) Final segmented images were used for crown root trait quantification using RhizoVision Explorer.

### Diameter thresholding can be used to remove stem tissue from root images

After examining the data obtained from analyzing crown root images from 22 genotypes in RhizoVision (**Fig. 3A**), it became apparent that the largest RootDiameter range (3000mm-above) corresponded to stem tissue in the images (**Fig. 3B**). Rather than trying to crop out stem from the root images, we instead decided to apply diameter thresholding to remove confounding stem data from the image analysis. After removing the highest RootDiameter range (3000mm-above) from the data, the resulting data was derived solely from root tissue (**Fig. 3C**). These data were then used for subsequent crown root trait quantification in RhizoVision.

**Figure 3.**
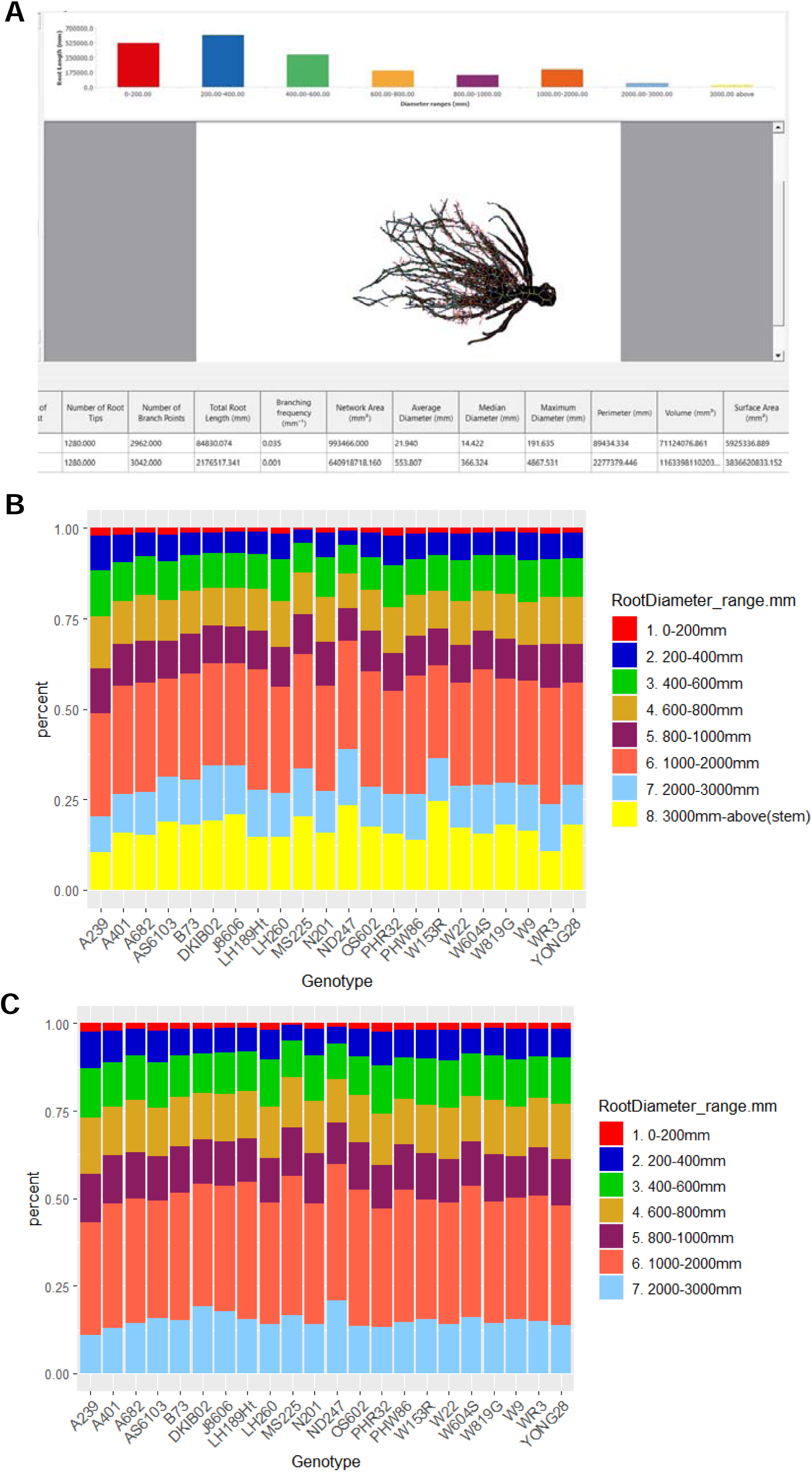
Diameter thresholding using RhizoVision to remove stem data. (A) Representative image from RhizoVision showing the root diameter thresholding output. The histogram indicates the distribution of total root lengths to each diameter range. (B) Root diameter ranges were measured in RhizoVision from maize crown root images across all genotypes sampled. The diameter range of 3000mm above (range 8) was the range found to correspond to the stem of the plant and not root tissue. (C) Root diameter ranges across all genotypes examined after removal of the 3000mm-above RootDiameter range category.

### Variability in Maize Crown Root Phenotypes

To determine the phenotypic range of crown root traits across maize inbreds, crown roots from 22 genotypes were analyzed using RhizoVision. In total, RhizoVision was able to extract 48 features from these crown root images. The trait values were examined relative to B73, the reference maize genotype, and clustered based on normalized Z-score (**Fig. 4**). Non-size related traits such as medium angle frequency and solidity group together on the left of the heatmap.

**Figure 4.**
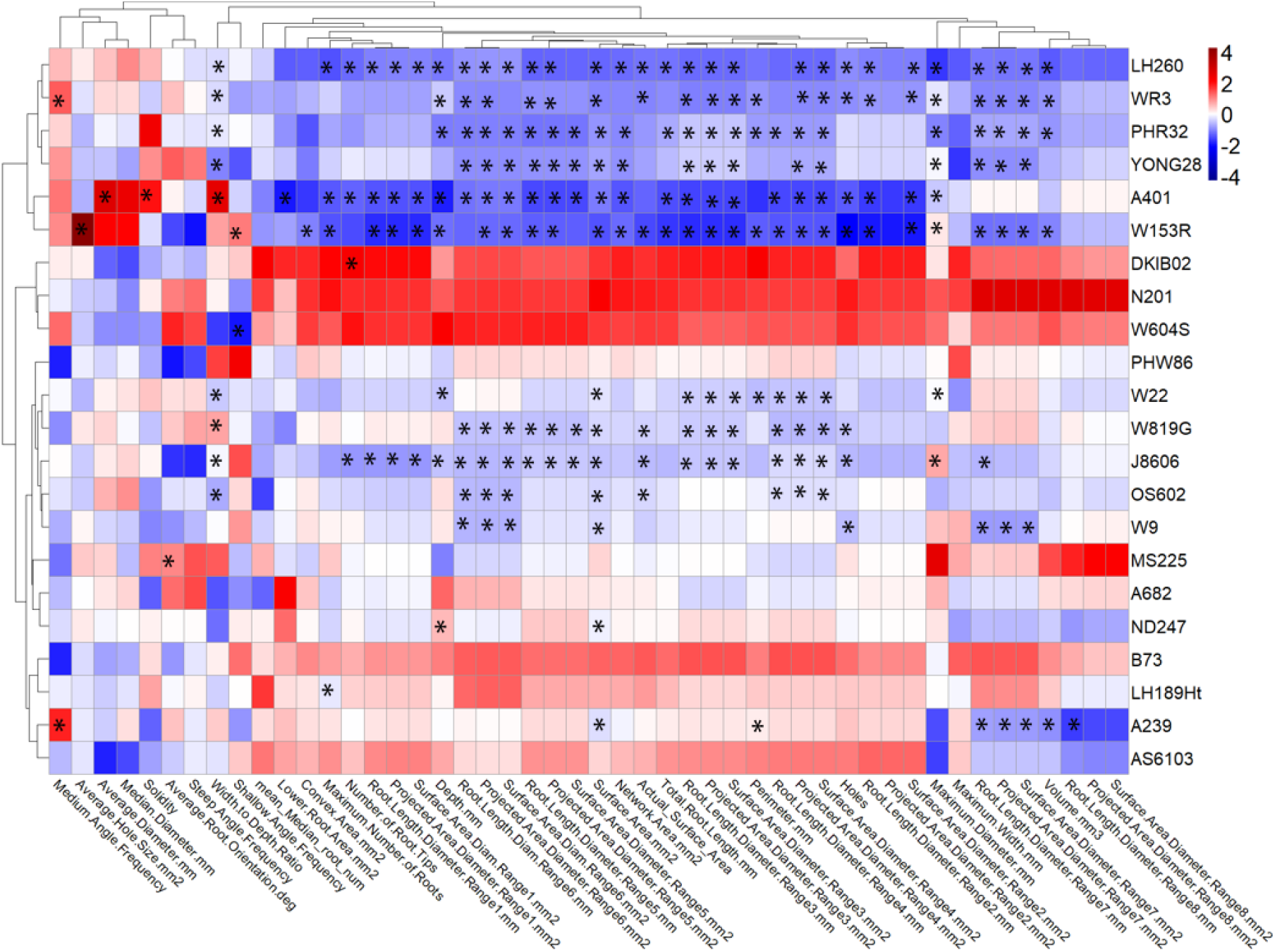
Hierarchical clustering of root traits across 22 genotypes. Quantification of 48 root traits (x-axis) in RhizoVision for 22 maize genotypes (y-axis). Normalized Z-scores for each trait are indicated on the heatmap compared to the mean value of B73 for each trait. The heatmap legend indicates the color in relationship to the Z-score. Statistical significance was assessed by a two-way analysis of variance (ANOVA) and indicated by an asterisk (*).

Notably, many genotypes exhibited different phenotypes in both size- and non-size-related traits. Overall, W153R contained the highest concentration of statistically significant phenotypes relative to B73. In contrast, PHW86, A682, and AS6103 exhibited similar root phenotypes as B73. Traits related to total root biomass, such as surface area and volume, were highly variable across the genotypes examined. Among size-related traits, several genotypes with smaller total root biomass were statistically significant from B73. This includes a cluster of 6 genotypes, including LH260, WR3, PHR32, YONG28, A401, and W153R (**Fig. 4**). Over half of the genotypes examined (13 out of 22) exhibited reduced surface area relative to B73 in these genotypes (Supplemental Table S6).

Non-size related traits such as medium.angle.frequency and solidity are grouped on the left of the heat map. These traits did not have a clear pattern of statistical significance among the genotypes examined. In addition, these non-size-related traits did not exhibit distinct clustering based on genotypes, suggesting a higher degree of variability in these root phenotypes.

### Gene expression patterns in maize roots with diverse phenotypes

To explore relationships between gene expression patterns and root phenotypes, we performed transcriptomic analysis on a subset of 11 inbreds that were phenotyped for crown root traits (**Fig. 4**). For this analysis, we examined transcript abundance relative to B73, the reference genotype for maize that displays median crown root phenotypes. All genotypes examined showed numerous differentially expressed (DE) transcripts relative to B73 in roots (**Table 1**). Depending on the genotype, the total number of differentially expressed genes (DEG) ranged from 140 to 4,477. In addition, most of the DEG were unique to a given genotype (Fig. 5) suggesting a strong genetic basis for these differences.

**Figure 5.**
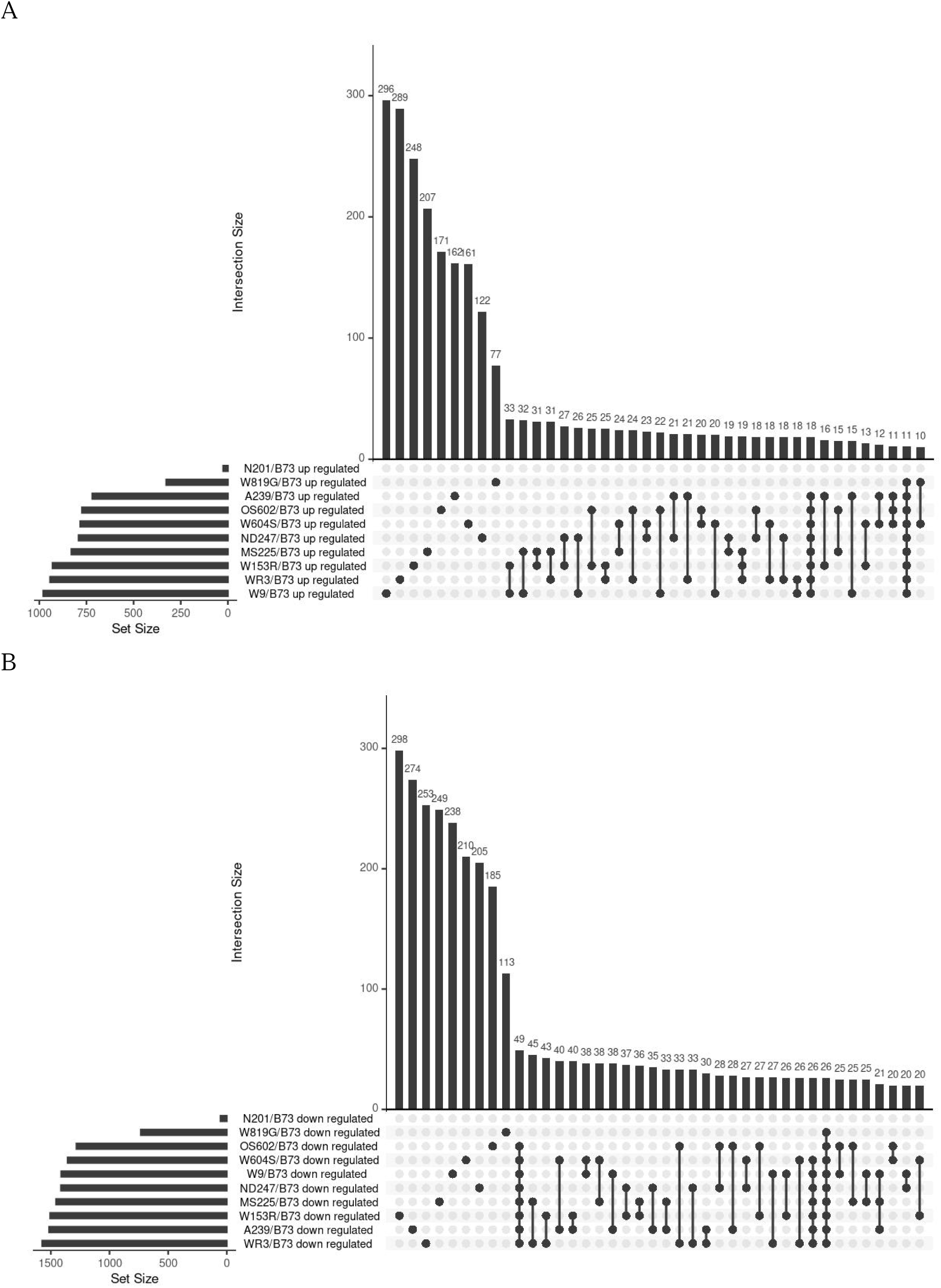
Overlap between differentially expressed genes among maize inbreds. **(A)** Upset plot of transcripts that were increased in maize inbreds relative to B73. **(B)** Upset plot of transcripts that were decreased in maize inbreds relative to B73.

We explored the overlap between DEG across genotypes using upset plots (**Fig. 5**). Most of the transcripts that displayed increased abundance relative to B73 were only present in one genotype (**Fig. 5A**). Very few transcripts with increased abundance were in common among al1 ten genotypes. A group of 11 transcripts was shared among nine genotypes in the upregulated datasets, which includes a gene associated with primary and secondary root elongation (*Zm00001d023213*), an essential root development enzyme (*Zm00001d005458*), and a gene connected to root cortex elongation (*Zm00001d037261*) (**Fig. 5A**). Among the transcripts that displayed reduced abundance among the ten genotypes relative to B73, most were unique to individual genotypes (**Fig. 5B**). A group of 26 transcripts was decreased in most genotypes examined, which includes genes associated with root development (e.g. *Zm00001d012955*, *Zm00001d040277*, and *Zm00001d040483*).

We also examined the top 100 DEG to see if any patterns emerged among these transcripts (**Fig. 6**). Notably, the ‘top 100’ DEG show genotype-specific differences in transcript abundance relative to B73. Specifically, all genotypes normalized to B73 show a difference in gene expression ND247, OS602, W153R, and WR3 have the most significant number of transcripts with increased abundance relative to B73, N201 exhibited the most significant degree of reduced transcript abundance compared to B73 (**Fig. 6**). There are four distinct upregulated and three downregulated clusters, GO Biological analysis (Ge, Jung, and Yao 2020; Tian et al. 2017; Du et al. 2010) performed on each cluster revealed several high-level categories of interest (The UniProt Consortium 2023) (Supplemental Table S8). Among the four upregulated groups found in W153R/B73, A239/B73, MS225/B73, ND247/B73, OS602/B73 and WR3/B73 all share cellular component organization (GO:0016043), cellular component organization or biogenesis (GO:0071840), establishment of localization (GO:0051234), localization (GO:0051179), protein folding (GO:0006457), response to chemical (GO:0042221), and stress response (GO:0006950) GO biology terms were found in each clustering group. Cellular component organization (GO:0016043) and cellular component organization or biogenesis (GO:0071840) are parent terms for cell wall organization (GO:0071555), which positively regulates positive regulation of cell wall organization or biogenesis (GO:1903340). Localization (GO:0051179) is a high-level parent term of auxin transport (GO:0060918). In cluster groups of A239/B73 and MS225/B73, W153R/B73, and WR3/B73 show between one and two genes associated with regulation of biological quality (GO:0065008) which is a closer parent term of hormone transport (GO:0009914) and auxin transport (GO:0060918). In W153R/B73 exhibits one gene associated with multicellular organism process (GO:0032501) and one gene associated with anatomical structure development (GO:0048856), parent term for primary, lateral, and root development (GO:0080022; GO:0048527; GO:0048364). W819G/B73 and N201/B73 established clusters of downregulated genes and all 100 genes in N201 were downregulated compared to B73, in addition to all the terms found in the upregulated groups, the most abundant groups were establishment of localization (GO:0051234) contained 13 genes, localization (GO:0051179) contained 13 genes, response to stimulus (GO:0050896) contained 12 genes, and stress response (GO:0006950) contained ten genes. Cell growth (GO:0016049), developmental growth (GO:0048589), signaling (GO:0023052), and response to chemicals (GO:0042221). W819G cellular component organization (GO:0071840), establishment of localization (GO:0051234), and localization (GO:0051179) contain four corresponding genes each. Among these ‘top 100’ DEG, none were observed across all genotypes, suggesting that genotype was a strong driver of the observed gene expression patterns in maize roots.

**Figure 6.**
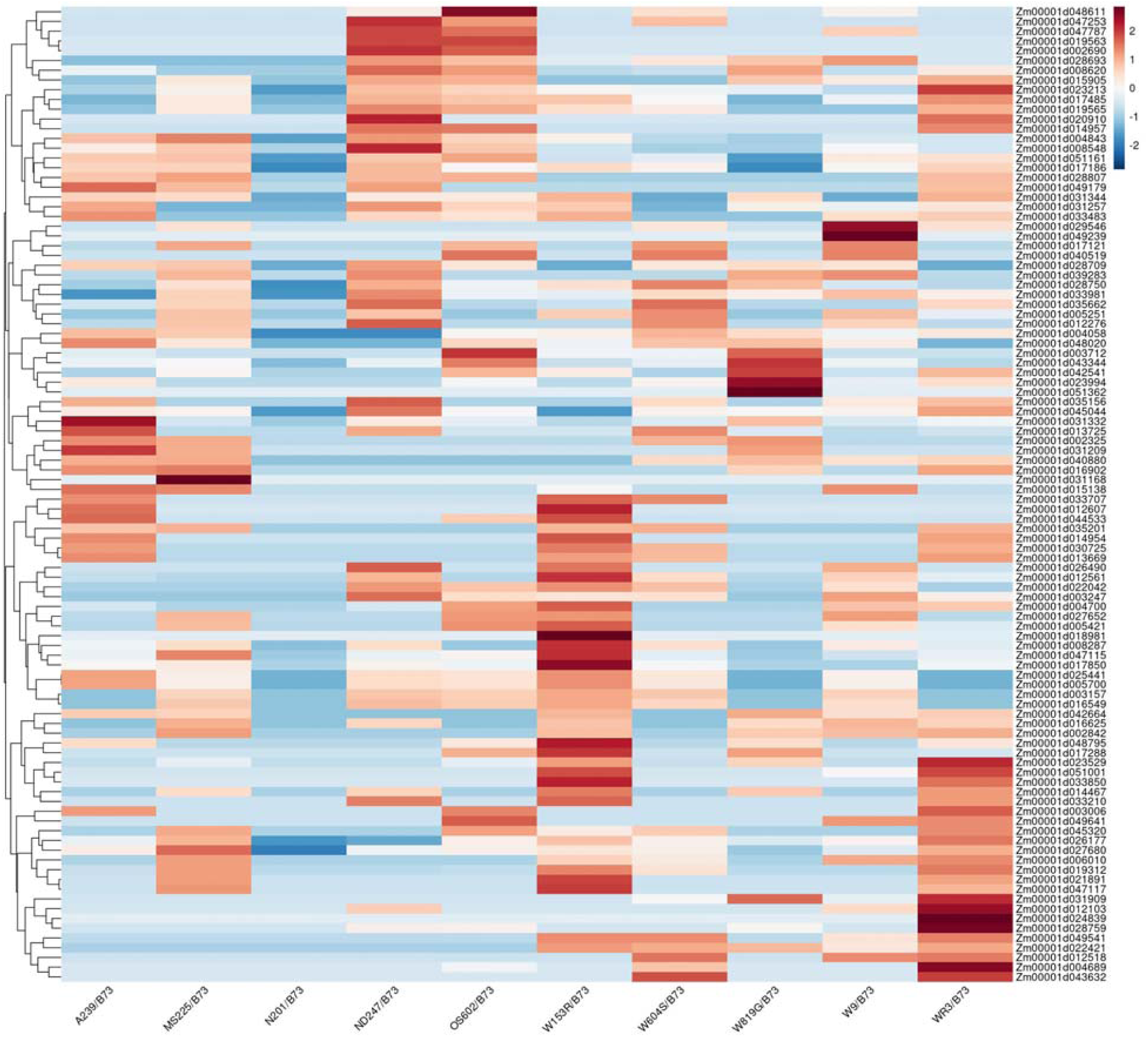
Hierarchical clustering of top 100 differentially expressed genes (DEGs). Clustering of the top 100 DEGs based on abundance in 10 different genotypes compared to B73. Scale bar indicates normalized transcript abundance relative to B73 per genotype.

### Maize root transcripts are enriched for hormone and stress GO terms

Molecular phenotypes among the DEG in maize roots across genotypes were examined by Gene Ontology (GO) enrichment analysis (**Supplemental Table S4**). We focused on GO biological process (GOBP) to make inferences about commonalities among the DEG. In total, 195 unique GO biological process (BP) terms were identified, with three terms found to be shared across all genotypes: secondary metabolic process (GO:0019748), single-organism metabolic process (GO:0044710), and small molecule metabolic process (GO:0044281) (**Fig. 7; Supplemental Table S4**). In addition, a subset of twenty terms related to root growth and development were of interest (**Fig. 7**). Across all genotypes, numerous terms related to stress response and cell wall categories are also enriched among the DEG in maize roots. Specifically, response to salt stress (GO:0009651), response to oxidative stress (GO:0006979), response to abiotic stimulus (GO:0009628), defense response to another organism (GO:0098542), and cell wall organization or biogenesis (GO:0071554) were all among the enriched GOBP. Because hormones are also key drivers of root development, other GOBP terms of interest included auxin metabolic process (GO:0009850), response to auxin (GO:0009733), response to jasmonic acid (GO:0009753), and response to karrikin (GO:0080167), and response to salicylic acid (GO:0009751) (**Fig. 7**).

**Figure 7.**
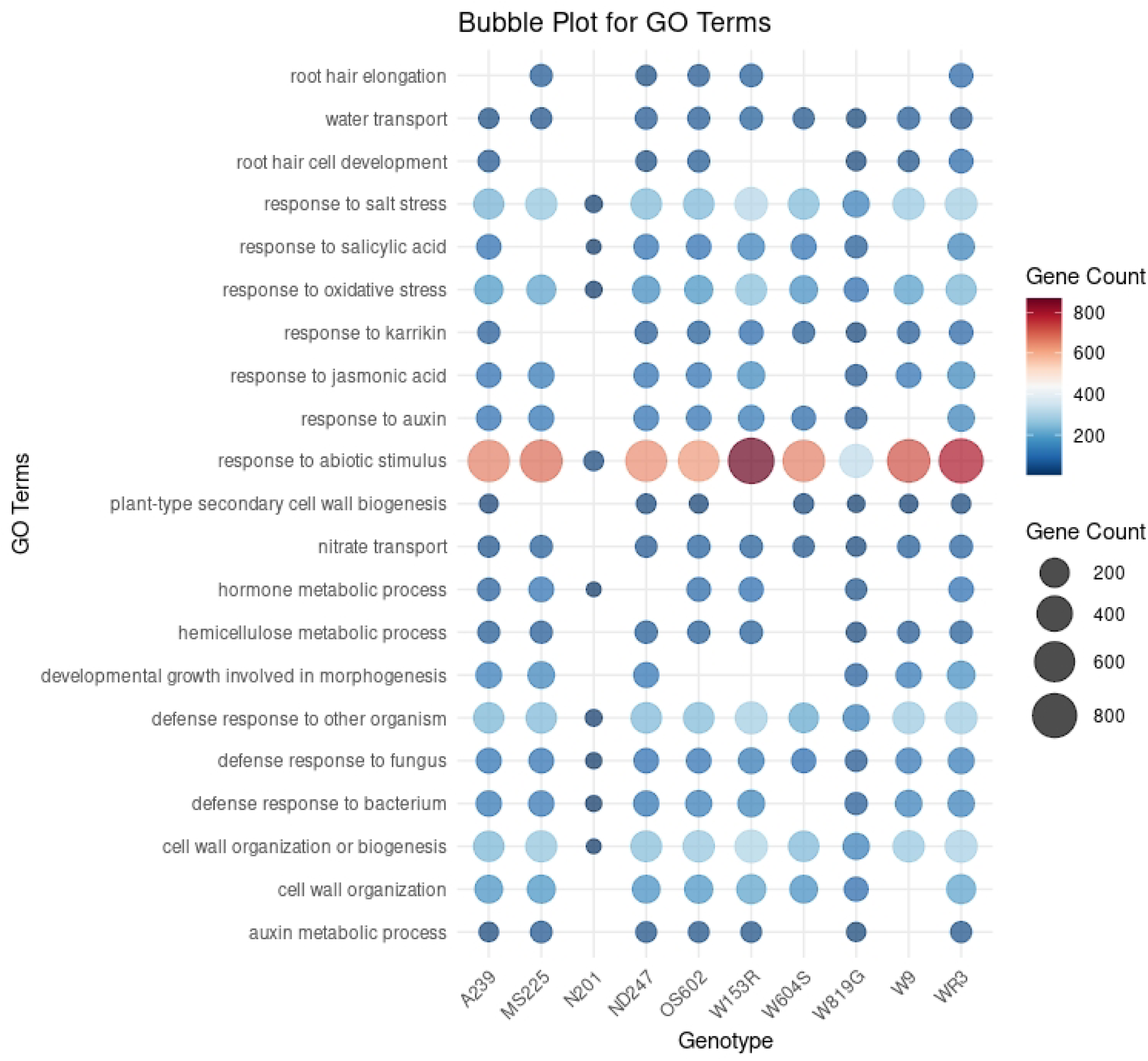
Enriched Gene Ontology (GO) terms among differentially expressed genes (DEG) in maize roots. GO terms related to biological process among DEG in maize roots across genotypes are indicated within each genotype as dots. The size and color of the dot correspond to the number of DEG underlying a given enriched term, as indicated in the legend.

### Juvenile and adult root traits are poorly correlated

Previous analyses have examined whether juvenile traits can predict adult traits in corn (Z. Liu et al. 2017; Leiboff et al. 2015). In both studies, the correlations between juvenile and adult phenotypes were inconsistent. We thus decided to leverage our datasets (Kelley et al. 2024; Kelley, Draves, and Muench 2024) to examine this relationship. To explore if juvenile maize root traits can be a predictor of adult root traits, we measured primary root length in 10-day-old (10DO) maize seedlings grown via rolled towel assays (Kelley, Draves, and Muench 2024) for 15 genotypes (5-10 seedlings per genotype) that had also been phenotyped in the field for crown root traits (**Fig. 8**). The correlation between primary root length in 10DO seedlings with 56 adult crown root traits was then calculated using Spearman correlation analysis (**Supplemental Table S5**). Among the 56 adult crown root traits analyzed for these genotypes, weak correlations (≥ 0.1-0.3) and moderate correlations (0.3-0.5) were observed among five genotypes (**Fig. 8**). This included A401, A682, AS6103, YONG28, and W9. Many of the genotypes examined (10/15) did not display any correlation between primary root length and adult root traits (**Fig. 8**). Overall, the correlation between primary root length and adult crown root traits appears to be genotype dependent.

**Figure 8.**
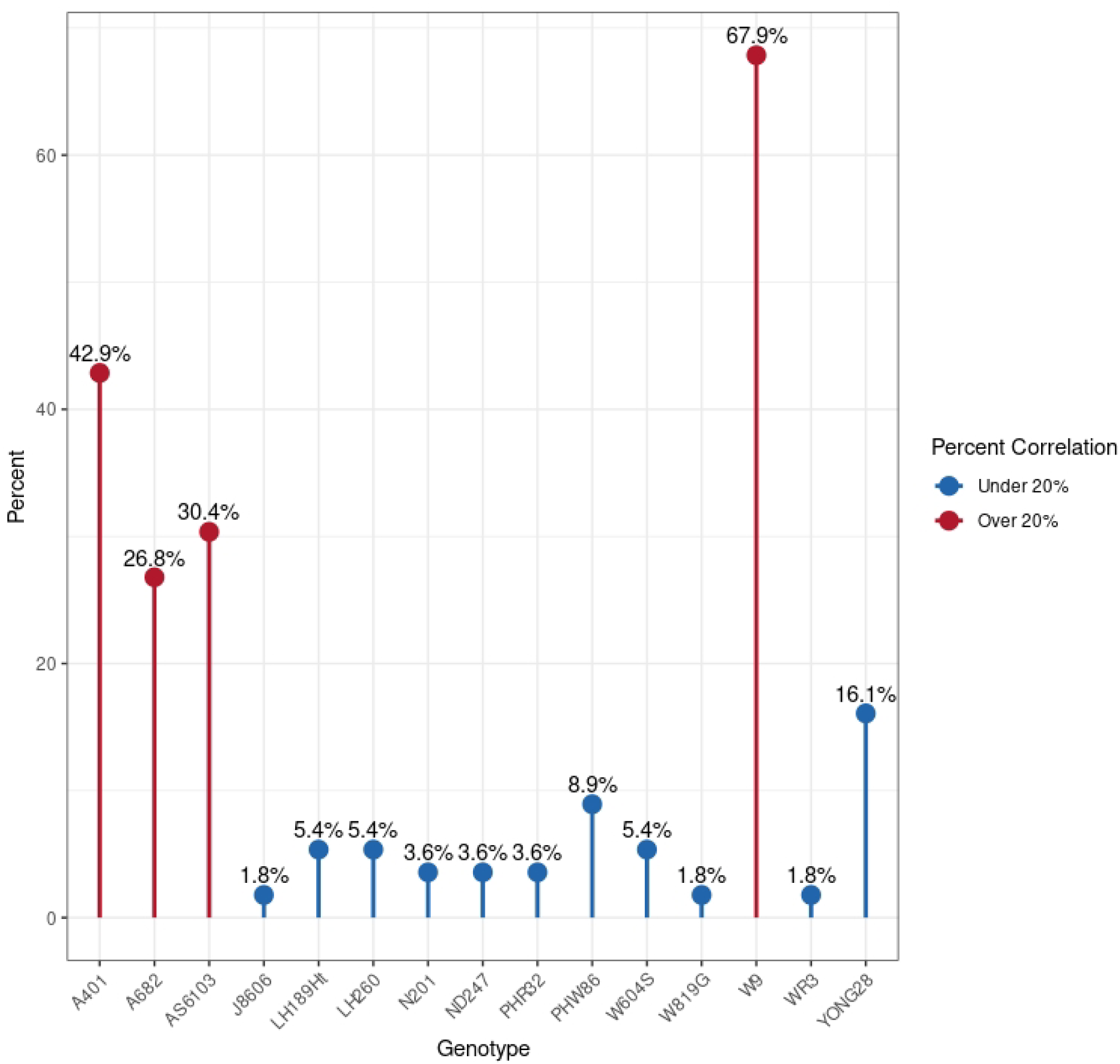
Spearman correlations between juvenile primary root length and adult crown root traits. The percentage of correlated phenotypes (y-axis) across 15 genotypes (x-axis) is indicated with a threshold of 20%.

## Discussion

This study focused on developing an affordable and feasible high throughput phenotypic extraction pipeline for maize roots to quantify crown root traits from field-grown plants. In addition, we examined transcriptomic signatures among several genotypes to examine genotype-phenotype relationships that may underpin root morphogenesis in corn. Finally, we leveraged juvenile root trait data to examine whether primary root length in seedlings can predict adult crown phenotypes. This study utilizes open-source software and includes publicly available root datasets relevant to the community.

Manual curation of plant phenotypes is laborious and time-consuming. A vital aspect of the image processing pipeline presented here is that it reduces user input quickly and efficiently. While our study does not compare a ground truth dataset to our pipeline outputs, previous work indicates that RhizoVision accurately measures root phenotypes (Seethepalli et al. 2021). Adult root phenotypes are complex to extract due to their complexity. Herein, we demonstrate that RootPainter can be trained on complex root image datasets to segment images effectively.

However, overlapping roots within an image are difficult to reconcile due to the complexity of 3D root systems being converted into 2D images. Thus, the background area(s) between these roots are still challenging for RootPainter to correctly segment. We leveraged ImageJ/Fiji to overcome this hurdle and properly differentiate such areas between fine roots. While this computational process slows pipeline processing, it dramatically reduces the time spent preparing each image. Finally, the segmented images are entered into RhizoVision for trait quantification (Seethepalli et al. 2020; 2021). In the future, this process could be automated for batch processing to increase throughput. While not completely free of human intervention, all images could be prepared, segmented, and phenotyped in approximately seven hours by one individual on one GPU. Due to the complexity of the roots, additional RootPainter training could result in a higher degree of segmentation in areas of overlapping roots. Overall, this workflow dramatically improves manual image measurement compared to hand-processing measurements with ImageJ/Fiji. It also is available to any researcher as RootPainter and RhizoVision are freely available open-source software. As a proof of concept, we applied this pipeline to 20 diverse maize genotypes grown in the field and observed a wide variation in crown root traits.

While the imaging pipeline presented here is highly transferable to other datasets, there are several limitations in the generation of training image sets including the need for human intervention in image preparation and GPU access. A substantial dataset of images with minimal noise is crucial for effectively training the model. Noise refers to additional objects in the image that could be mistaken for structures of interest. In this case, that refers to moisture and dirt miming the roots profile, requiring additional image cleaning. Images require additional processing to reduce noise and enable RhisoVision to extract root features accurately. Despite advancements, user input is necessary to prepare images for feature extraction. Due to the complexity of the root systems and the quality of images, additional training beyond the 128 initial images could increase the accuracy of the segmented images. The presented here model is currently limited to maize adult crown root images without stem tissue. Additional instances of RootPainter can be trained to extract traits from plants of various species and development stages. GPU access through Google Colab (Bisong 2019) could provide the infrastructure necessary to retrain RootPainter, allowing different individuals to work on this process regardless of location or VPN capabilities.

Morphological phenotypes can be driven by molecular phenotypes such as gene expression. To examine how transcriptome patterns vary across maize genotypes with diverse crown root phenotypes, we performed QuantSeq 3’mRNA analysis on seedling roots. In total, thousands of transcripts showed variable expression levels relative to B73, the reference genotype for maize. Notably, very few individual transcripts were shared among the genotypes examined relative to B73. However, GO enrichment analysis of biological process terms revealed striking similarities among DEG across several genotypes. This includes terms logical for root physiology, such as abiotic stress and hormone pathways. Thus, while the unique gene products may not be shared across genotypes their associated annotated biological pathways are maintained. Overall, the transcriptome analysis suggests that hormones such as auxin, cell wall biogenesis, and stress responses may impact maize crown root phenotypes. These data provide an excellent resource for future investigation into examining these gene products individually and/or collectively to better understand which transcripts influence certain aspects of maize root development.

Adult field-grown phenotyping is time and labor-intensive. There has been discussion about whether seedling phenotype can accurately predict adult phenotypes to reduce the amount of material that should be examined in breeding pipelines. However, very few studies have integrated juvenile data with adult data to address the feasibility of this idea. Two studies previously found inconsistent correlations between juvenile and adult maize phenotypes (Pace et al. 2015). Here, we observed the same result for most, but not all, of the genotypes we examined. In comparing juvenile and adult root traits in 15 maize genotypes, we observed only five genotypes with a weakly to moderate positive correlation between 10-day-old primary root length and adult crown root traits. While these findings align with previous research, it has yet to be determined why a subset of genotypes express correlations in phenotypes between developmental stages. In addition, these results suggest that relationships between juvenile and adult phenotypes are complex and may be heavily influenced by genotype. Additional investigation into the plasticity and genetic basis of maize root development will be informative for better understanding the inconsistent correlations between seedling and adult phenotypes. Notably, W9 exhibited a high correlation between 10DO primary root length and several adult crown root phenotypes. In the GO analysis of DEG in W9 relative to B73, we observed several missing GO term enrichments relative to the other genotypes examined, including root hair elongation, response to auxin, hormone metabolic process, cell wall organization, and auxin metabolic process. Auxin plays a pivotal role in maize root growth (Y. Zhang et al. 2015; Kong et al. 2024; Wang et al. 2023; M. Zhang et al. 2018; Zheng et al. 2023; J. Li et al. 2022; Cowling, Dash, and Kelley 2023; Y. Zhang et al. 2014; Cowling et al. 2024; Z. Li et al. 2018). Thus, the observed lack of auxin pathway transcripts enriched in W9 roots could underpin the phenotypic constraint between seedling and adult states. Auxin promotes rapid growth in cell expansion (Du, Spalding, and Gray 2020), and a distinct lack of enrichment in auxin-specific biological processes could contribute to the observed proportional growth in W9.

In conclusion, the RootPainter to RhizoVision pipeline employed here provides an effective high-throughput root phenotype quantification workflow. This pipeline enabled analysis of juvenile and adult root phenotype correlations, exploration of genotype-phenotype relationships, and open dataset generation. Through this work we have outlined an effective method of root phenotype quantification that can be used by any researcher with free open-source software. This study can inform future studies into the role of hormones such as auxin in shaping diverse maize root phenotypes, the genetic basis of root phenotypic plasticity, and refined approaches for high-throughput root phenotyping.

## Acknowledgments

This work was supported by the United States Department of Agriculture (USDA) National Institute of Food and Agriculture (NIFA) Agriculture and Food Research Initiative (AFRI) award number GRANT12907916 to DRK and JWW; Hatch Act State of Iowa funds IOW03649 and IOW05745 to DRK; Hatch Act and State of Iowa funds IOW04108 to JWW; the Iowa State University Plant Science Institute (JWW); the ISU Crop Bioengineering Center (DRK and JWW); and a Department of Defense (DOD) Science, Mathematics, and Research for Transformation (SMART) scholarship to JBC. HS was supported by NSF BIORETS award number 2147083. Thank you to Yitzhe Tang for assistance with training RootPainter.

## Data Availability

All data, code, and genetic materials used in this study are available to any researcher for purposes of reproducing or extending the analysis. Maize crown root image data is available at DataShare, the Iowa State University Open Research Data Repository (Kelley et al. 2024). The maize seedling image dataset is available at DataShare, the Iowa State University Open Research Data Repository (Kelley, Draves, and Muench 2024). Transcriptomic data is deposited at the Sequence Read Archive (SRA) as submission SUB# PRJNA1108420. All other data are available in the main text or the supplementary materials.

## References

Amtmann, Anna, Malcolm J. Bennett, and Amelia Henry. 2022. “Root Phenotypes for the Future.” Plant, Cell & Environment 45 (3): 595–601. 10.1111/pce.14269.

Armengaud, Patrick, Kevin Zambaux, Adrian Hills, Ronan Sulpice, Richard J. Pattison, Michael R. Blatt, and Anna Amtmann. 2009. “EZ-Rhizo: Integrated Software for the Fast and Accurate Measurement of Root System Architecture.” Plant Journal 57 (5): 945–56. 10.1111/j.1365-313X.2008.03739.x.

Bauer, Felix Maximilian, Lena Lärm, Shehan Morandage, Guillaume Lobet, Jan Vanderborght, Harry Vereecken, and Andrea Schnepf. 2022. “Development and Validation of a Deep Learning Based Automated Minirhizotron Image Analysis Pipeline.” Plant Phenomics 2022 (May):9758532. 10.34133/2022/9758532.

Cai, Jinhai, Zhanghui Zeng, Jason N. Connor, Chun Yuan Huang, Vanessa Melino, Pankaj Kumar, and Stanley J. Miklavcic. 2015. “RootGraph: A Graphic Optimization Tool for Automated Image Analysis of Plant Roots.” Journal of Experimental Botany 66 (21): 6551–62. 10.1093/jxb/erv359.

Chen, Ying Long, Ivica Djalovic, and Zed Rengel. 2015. “Phenotyping for Root Traits.” In Phenomics in Crop Plants: Trends, Options and Limitations, edited by Jitendra Kumar, Aditya Pratap, and Shiv Kumar, 101–28. New Delhi: Springer India. 10.1007/978-81-322-2226-2_8.

Cowling, Craig L, Linkan Dash, and Dior R Kelley. 2023. “Roles of Auxin Pathways in Maize Biology.” Journal of Experimental Botany 74 (22): 6989–99. 10.1093/jxb/erad297.

Cowling, Craig L., Arielle L. Homayouni, Jodi B. Callwood, Maxwell R. McReynolds, Jasper Khor, Haiyan Ke, Melissa A. Draves, et al. 2024. “ZmPILS6 Is an Auxin Efflux Carrier Required for Maize Root Morphogenesis.” Proceedings of the National Academy of Sciences of the United States of America 121 (22): e2313216121. 10.1073/pnas.2313216121.

Das, Abhiram, Hannah Schneider, James Burridge, Ana Karine Martinez Ascanio, Tobias Wojciechowski, Christopher N. Topp, Jonathan P. Lynch, Joshua S. Weitz, and Alexander Bucksch. 2015. “Digital Imaging of Root Traits (DIRT): A High-Throughput Computing and Collaboration Platform for Field-Based Root Phenomics.” Plant Methods 11 (November):51. 10.1186/s13007-015-0093-3.

Draves, Melissa A., Rebekah L. Muench, Michelle G. Lang, and Dior R. Kelley. 2022. “Maize Seedling Growth and Hormone Response Assays Using the Rolled Towel Method.” Current Protocols 2 (10): e562. 10.1002/cpz1.562.

Du, Zhou, Xin Zhou, Yi Ling, Zhenhai Zhang, and Zhen Su. 2010. “agriGO: A GO Analysis Toolkit for the Agricultural Community.” Nucleic Acids Research 38 (suppl_2): W64–70. 10.1093/nar/gkq310.

French, Andrew, Susana Ubeda-Tomás, Tara J. Holman, Malcolm J. Bennett, and Tony Pridmore. 2009. “High-Throughput Quantification of Root Growth Using a Novel Image-Analysis Tool.” Plant Physiology 150 (4): 1784–95. 10.1104/pp.109.140558.

Gage, Joseph L., Nathan D. Miller, Edgar P. Spalding, Shawn M. Kaeppler, and Natalia de Leon. 2017. “TIPS: A System for Automated Image-Based Phenotyping of Maize Tassels.” Plant Methods 13 (1): 1–12. 10.1186/s13007-017-0172-8.

Gärtner, Holger, Bettina Wagner, Ingo Heinrich, and Clemens Denier. 2009. “3D-Laser Scanning: A New Method to Analyze Coarse Tree Root Systems.” For. Snow Landsc. Res 82 (January):95–106.

Ge, Steven Xijin, Dongmin Jung, and Runan Yao. 2020. “ShinyGO: A Graphical Gene-Set Enrichment Tool for Animals and Plants.” Bioinformatics 36 (8): 2628–29. 10.1093/bioinformatics/btz931.

Gehan, Malia A., Noah Fahlgren, Arash Abbasi, Jeffrey C. Berry, Steven T. Callen, Leonardo Chavez, Andrew N. Doust, et al. 2017. “PlantCV v2: Image Analysis Software for High-Throughput Plant Phenotyping.” PeerJ 2017 (12): e4088–e4088. 10.7717/peerj.4088.

Guo, Li, Jin Chen, Xihong Cui, Bihang Fan, and Henry Lin. 2013. “Application of Ground Penetrating Radar for Coarse Root Detection and Quantification: A Review.” Plant and Soil 362 (1): 1–23. 10.1007/s11104-012-1455-5.

He, Kunhui, Zheng Zhao, Wei Ren, Zhe Chen, Limei Chen, Fanjun Chen, Guohua Mi, Qingchun Pan, and Lixing Yuan. 2023. “Mining Genes Regulating Root System Architecture in Maize Based on Data Integration Analysis.” Theoretical and Applied Genetics 136 (6): 127. 10.1007/s00122-023-04376-0.

Hochholdinger, Frank, Peng Yu, and Caroline Marcon. 2018. “Genetic Control of Root System Development in Maize.” Trends in Plant Science 23 (1): 79–88. 10.1016/j.tplants.2017.10.004.

Kelley, Dior, Melissa Draves, Jackson Marshall, Craig Cowling, and Heather Sweers. 2024. “Maize Crown Root Images from 22 Genotypes,” April. 10.25380/iastate.25541068.v1.

Kelley, Dior, Melissa Draves, and Rebekah Muench. 2024. “Maize Seedling Images from 617 Genotypes.” https://iastate.figshare.com/: Iowa State University. 10.25380/iastate.25625430.v1.

Kong, Xiuzhen, Yali Xiong, Xiaoyun Song, Samuel Wadey, Suhang Yu, Jinliang Rao, Aneesh Lale, et al. 2024. “Ethylene Regulates Auxin-Mediated Root Gravitropic Machinery and Controls Root Angle in Cereal Crops.” Plant Physiology 195 (3): 1969–80. 10.1093/plphys/kiae134.

Leiboff, Samuel, Xianran Li, Heng-Cheng Hu, Natalie Todt, Jinliang Yang, Xiao Li, Xiaoqing Yu, et al. 2015. “Genetic Control of Morphometric Diversity in the Maize Shoot Apical Meristem.” Nature Communications 6 (November):8974. 10.1038/ncomms9974.

Li, Jing, Fengkai Wu, Yafeng He, Bing He, Ying Gong, Baba Salifu Yahaya, Yuxin Xie, et al. 2022. “Maize Transcription Factor ZmARF4 Confers Phosphorus Tolerance by Promoting Root Morphological Development.” International Journal of Molecular Sciences 23 (4): 2361. 10.3390/ijms23042361.

Li, Pengcheng, Zhihai Zhang, Gui Xiao, Zheng Zhao, Kunhui He, Xiaohong Yang, Qingchun Pan, et al. 2024. “Genomic Basis Determining Root System Architecture in Maize.” Theoretical and Applied Genetics 137 (5): 102. 10.1007/s00122-024-04606-z.

Li, Zhaoxia, Xinrui Zhang, Yajie Zhao, Yujie Li, Guangfeng Zhang, Zhenghua Peng, and Juren Zhang. 2018. “Enhancing Auxin Accumulation in Maize Root Tips Improves Root Growth and Dwarfs Plant Height.” Plant Biotechnology Journal 16 (1): 86–99. 10.1111/pbi.12751.

Liu, Suxing, Carlos Sherard Barrow, Meredith Hanlon, Jonathan P. Lynch, and Alexander Bucksch. 2021. “DIRT/3D: 3D Root Phenotyping for Field-Grown Maize (Zea Mays).” Plant Physiology 187 (2): 739–57. 10.1093/plphys/kiab311.

Liu, Zhigang, Kun Gao, Shengchen Shan, Riling Gu, Zhangkui Wang, Eric J. Craft, Guohua Mi, Lixing Yuan, and Fanjun Chen. 2017. “Comparative Analysis of Root Traits and the Associated QTLs for Maize Seedlings Grown in Paper Roll, Hydroponics and Vermiculite Culture System.” Frontiers in Plant Science 8:436. 10.3389/fpls.2017.00436.

Love, Michael I., Simon Anders, Vladislav Kim, and Wolfgang Huber. 2015. “RNA-Seq Workflow: Gene-Level Exploratory Analysis and Differential Expression.” F1000Research. 10.12688/f1000research.7035.1.

Mahesh Kumar, K., and A. Rama Mohan Reddy. 2016. “A Fast DBSCAN Clustering Algorithm by Accelerating Neighbor Searching Using Groups Method.” Pattern Recognition 58 (October):39–48. 10.1016/j.patcog.2016.03.008.

Mohamed, Awaz, Yogan Monnier, Zhun Mao, Guillaume Lobet, Jean-Luc Maeght, Merlin Ramel, and Alexia Stokes. 2017. “An Evaluation of Inexpensive Methods for Root Image Acquisition When Using Rhizotrons.” Plant Methods 13 (1): 11. 10.1186/s13007-017-0160-z.

Pace, Jordon, Candice Gardner, Cinta Romay, Baskar Ganapathysubramanian, and Thomas Lübberstedt. 2015. “Genome-Wide Association Analysis of Seedling Root Development in Maize (Zea Mays L.).” BMC Genomics 16 (1): 47. 10.1186/s12864-015-1226-9.

Pace, Jordon, Nigel Lee, Hsiang Sing Naik, Baskar Ganapathysubramanian, and Thomas Lübberstedt. 2014. “Analysis of Maize (Zea Mays L.) Seedling Roots with the High-Throughput Image Analysis Tool ARIA (Automatic Root Image Analysis).” Edited by Pawan L. Kulwal. PLoS ONE 9 (9): e108255–e108255. 10.1371/journal.pone.0108255.

Paez-Garcia, Ana, Christy M. Motes, Wolf-Rüdiger Scheible, Rujin Chen, Elison B. Blancaflor, and Maria J. Monteros. 2015. “Root Traits and Phenotyping Strategies for Plant Improvement.” Plants 4 (2): 334–55. 10.3390/plants4020334.

Pound, Michael P., Andrew P. French, Jonathan A. Atkinson, Darren M. Wells, Malcolm J. Bennett, and Tony Pridmore. 2013. “RootNav: Navigating Images of Complex Root Architectures.” Plant Physiology 162 (4): 1802–14. 10.1104/pp.113.221531.

Schauberger, Philipp, and Alexander Walker. 2022. Openxlsx: Read, Write and Edit Xlsx Files.

Schmidt, Jennifer E., Carolyn Lowry, and Amelie C.M. Gaudin. 2018. “An Optimized Rhizobox Protocol to Visualize Root Growth and Responsiveness to Localized Nutrients.” Journal of Visualized Experiments : JoVE, no. 140 (October), 58674. 10.3791/58674.

Schneider, Caroline A., Wayne S. Rasband, and Kevin W. Eliceiri. 2012. “NIH Image to ImageJ: 25 Years of Image Analysis.” Nature Methods 9 (7): 671–75. 10.1038/nmeth.2089.

Seethepalli, Anand, Kundan Dhakal, Marcus Griffiths, Haichao Guo, Gregoire T Freschet, and Larry M York. 2021a. “RhizoVision Explorer: Open-Source Software for Root Image Analysis and Measurement Standardization.” AoB PLANTS 13 (6): plab056. 10.1093/aobpla/plab056.

Seethepalli, Anand, Kundan Dhakal, Marcus Griffiths, Haichao Guo, Gregoire T Freschet, and Larry M York. 2021b. “RhizoVision Explorer: Open-Source Software for Root Image Analysis and Measurement Standardization.” AoB PLANTS 13 (6): plab056. 10.1093/aobpla/plab056.

Sha, Xiao-qian, Hong-hui Guan, Yu-qian Zhou, Er-hu Su, Jian Guo, Yong-xiang Li, Deng-feng Zhang, et al. 2023. “Genetic Dissection of Crown Root Traits and Their Relationships with Aboveground Agronomic Traits in Maize.” Journal of Integrative Agriculture 22 (11): 3394–3407. 10.1016/j.jia.2023.04.022.

Smith, Abraham George, Eusun Han, Jens Petersen, Niels Alvin, Faircloth Olsen, Christian Giese, Miriam Athmann, Dorte Bodin Dresbøll, and Kristian Thorup-Kristensen. 2022. “RootPainter: Deep Learning Segmentation of Biological Images with Corrective Annotation” 236 (2): 774–91. 10.1101/2020.04.16.044461.

Steudle, Ernst. 2000. “Water Uptake by Plant Roots: An Integration of Views.” Plant and Soil 226 (1): 45–56. 10.1023/A:1026439226716.

The UniProt Consortium. 2023. “UniProt: The Universal Protein Knowledgebase in 2023.” Nucleic Acids Research 51 (D1): D523–31. 10.1093/nar/gkac1052.

Tian, Tian, Yue Liu, Hengyu Yan, Qi You, Xin Yi, Zhou Du, Wenying Xu, and Zhen Su. 2017. “agriGO v2.0: A GO Analysis Toolkit for the Agricultural Community, 2017 Update.” Nucleic Acids Research 45 (W1): W122–29. 10.1093/nar/gkx382.

Vogt, Kristiina A., Daniel J. Vogt, Heidi Asbjornsen, and Randy A. Dahlgren. 1995. “Roots, Nutrients and Their Relationship to Spatial Patterns.” Plant and Soil 168 (1): 113–23. 10.1007/BF00029320.

Wang, Yubin, Jiapeng Xing, Jiachi Wan, Qingqing Yao, Yushi Zhang, Guohua Mi, Limei Chen, Zhaohu Li, and Mingcai Zhang. 2023. “Auxin Efflux Carrier ZmPIN1a Modulates Auxin Reallocation Involved in Nitrate-Mediated Root Formation.” BMC Plant Biology 23 (1): 74. 10.1186/s12870-023-04087-0.

Wickham, Hadley, Mara Averick, Jennifer Bryan, Winston Chang, Lucy D’Agostino McGowan, Romain François, Garrett Grolemund, et al. 2019. “Welcome to the Tidyverse.” Journal of Open Source Software 4 (43): 1686. 10.21105/joss.01686.

Wickham, Hadley, Romain François, Lionel Henry, Kirill Müller, and Davis Vaughan. 2023. Dplyr: A Grammar of Data Manipulation. https://dplyr.tidyverse.org.

Yasrab, Robail, Jonathan A Atkinson, Darren M Wells, Andrew P French, Tony P Pridmore, and Michael P Pound. 2019. “RootNav 2.0: Deep Learning for Automatic Navigation of Complex Plant Root Architectures.” GigaScience 8 (giz123). 10.1093/gigascience/giz123.

York, Larry M., and Jonathan P. Lynch. 2015. “Intensive Field Phenotyping of Maize (Zea Mays L.) Root Crowns Identifies Phenes and Phene Integration Associated with Plant Growth and Nitrogen Acquisition.” Journal of Experimental Botany 66 (18): 5493–5505. 10.1093/jxb/erv241.

Zhang, Maolin, Xiaoduo Lu, Cuiling Li, Bing Zhang, Chunyi Zhang, Xian-sheng Zhang, and Zhaojun Ding. 2018. “Auxin Efflux Carrier ZmPGP1 Mediates Root Growth Inhibition under Aluminum Stress.” Plant Physiology 177 (2): 819–32. 10.1104/pp.17.01379.

Zhang, Yanxiang, Inga von Behrens, Roman Zimmermann, Yvonne Ludwig, Stefan Hey, and Frank Hochholdinger. 2015. “LATERAL ROOT PRIMORDIA 1 of Maize Acts as a Transcriptional Activator in Auxin Signalling Downstream of the Aux/IAA Gene Rootless with Undetectable Meristem 1.” Journal of Experimental Botany 66 (13): 3855–63. 10.1093/jxb/erv187.

Zhang, Yanxiang, Anja Paschold, Caroline Marcon, Sanzhen Liu, Huanhuan Tai, Josefine Nestler, Cheng-Ting Yeh, et al. 2014. “The Aux/IAA Gene Rum1 Involved in Seminal and Lateral Root Formation Controls Vascular Patterning in Maize (Zea Mays L.) Primary Roots.” Journal of Experimental Botany 65 (17): 4919–30. 10.1093/jxb/eru249.

Zheng, Zhigang, Baobao Wang, Chuyun Zhuo, Yurong Xie, Xiaoming Zhang, Yanjun Liu, Guisen Zhang, et al. 2023. “Local Auxin Biosynthesis Regulates Brace Root Angle and Lodging Resistance in Maize.” New Phytologist 238 (1): 142–54. 10.1111/nph.18733.

